# Population of the Paraná Antwren *Formicivora acutirostris* reassessed as being 64% smaller: Revision of its conservation status and assessment of its Green Status, with management proposals

**DOI:** 10.1101/2023.10.10.561509

**Authors:** Marcos R. Bornschein, Giovanna Sandretti-Silva, Daiane D. Sobotka, Leandro Corrêa, Bianca L. Reinert, Fabio Stucchi Vannucchi, Marcio R. Pie

## Abstract

Assessing the conservation status of a species is important for designing effective conservation measures. Consequently, it is often vital to review it to update biodiversity management. *Formicivora acutirostris* is a bird from coastal marshes and related habitats of Brazil’s southern flood plains. It is considered endangered in Brazil but near threatened globally. In 2007, its area of occupancy (AOO) and population size were estimated based on assessment of aerial photographs from 1978 and 1980. Since the species is threatened and occupies a small area scattered across a coastal pressured region, we aimed to reassess its conservation status and assess its Green Status. We compiled new records, conducted new density estimates, and compared the previous mapping with orbital imagery to estimate the current AOO, habitat loss, population size, and review its conservation status based on the International Union for Conservation of Nature criteria. The species is distributed across 10 populations, including two new populations from the southern coast of Santa Catarina to the northern coast of Rio Grande do Sul. We estimated the AOO at 4,102 ha and the population as 6,284 mature territorial individuals. The previously mapped AOO has decreased by 1,535 ha due to ecological succession. The loss of habitat due to invasion by exotic grasses is the main anthropogenic impact. We recommend that the species be considered vulnerable. The Green Status indicates that the Conservation Legacy of actions taken thus far and the Conservation Dependence of ongoing actions are null, but it highlights the importance of future actions for species conservation. We propose the establishment of exotic-free zones as small geographic areas with a significant amount of minimally invaded environments, which we suggest as priority areas for the conservation of the species due to their cost-effective management potential. We also propose assisted colonization to enhance its long-term conservation.

## Introduction

The geographical distribution and dynamics of a species are among its fundamental characteristics, as well as essential tools for assessing its extinction risk (Pearson 2007, Gaston and Fuller 2009, Syfert *et al*. 2014). The distribution of a species can be mapped simply by establishing its extreme sites of occurrence with large boundaries that encompass all environments within that polygon to determine the extent of occurrence (EOO). A more complex method includes evaluation of only the habitats actually occupied by the species to determine the area of occupancy (AOO; BirdLife International 2000, Gaston and Fuller 2009, International Union for Conservation of Nature [IUCN] 2012, Jiménez-Alfaro *et al*. 2012, IUCN 2022). The IUCN uses both methods as criteria to assess the conservation status of a species (IUCN 2012, 2022) because they express possible threats and vulnerabilities and are efficient for estimating population sizes (IUCN 2012, 2022). The population size of a species is another criterion used to classify its conservation status (IUCN 2012) and is more precise and appropriate when calculated according to the AOO (Gaston and Fuller 2009).

The Paraná Antwren *Formicivora acutirostris* is a species endemic to the Atlantic Forest biome (Brooks *et al*. 1999) in Southern Brazil, which was first described in 1995 (Bornschein *et al*. 1995). This species and its sister species, the Marsh Antwren *F. paludicola*, described in 2014, are the only marsh-dwelling thamnophilids worldwide (Zimmer and Isler 2003: 492, Buzzetti *et al*. 2013). The geographic distribution of *F. acutirostris* was proposed to be divided into eight isolated populations distributed from the central coast of Paraná to the north coast of Santa Catarina in Southern Brazil, totaling an estimated AOO of 6,060 ha and an estimated population of 17,680 mature individuals (Reinert *et al*. 2007).

*Formicivora acutirostris* has been assessed as near threatened (NT) globally (BirdLife International [2019]) and vulnerable (VU) in Brazil (Ordinance MMA no. 148/2022). The conservation of this species is a constant concern because it occupies a coastal region under pressure from real estate development, and also because its habitat is decreasing in area and quality due to agricultural expansion and the invasion of alien plant species (Reinert *et al*. 2007). Hence, the geographic distribution of the species assessed by Reinert *et al*. (2007) is currently outdated because it was based on aerial photographs taken over 40 years ago (in 1978 and 1980). In addition, the species was recorded in Rio Grande do Sul (Bencke *et al*. 2010), 329 km south of Reinert *et al*.’s (2007) southernmost site, suggesting a more positive conservation scenario. In the present study, we updated the estimates of the AOO and population size of *F. acutirostris* and reassessed its conservation status, as suggested previously (Reinert and Bornschein 2008, Reinert *et al*. 2009). Herein, we also propose potential conservation measures.

## Methods

### Target species

*Formicivora acutirostris* is a long-lived species (Bornschein *et al*. 2015) that forms permanent pairs inhabiting territories that are defended over time (Reinert 2008). The species displays sexual dimorphism (Reinert and Bornschein 1996) and has low flight capacity, typically not flying more than 25 m above vegetation (Reinert *et al*. 2007).

### Estimates of individual density

We worked at four sites in Guaratuba Bay, Ramsar Site Guaratuba, located in municipality of Guaratuba, on the Paraná coast, southern Brazil: Jundiaquara Island (*c.* 25°52’25”S, 48°45’32”W; 11.3 ha; Figure 1), the confluence of the Claro and São João Rivers (“Continente”; *c.* 25°52’28”S, 48°45’44”W; 8.5 ha; Figure 1), Folharada Island (*c.* 25°51’58”S, 48°43’23”W; 16.3 ha) and Lagoa do Parado (*c.* 25°44’36”S, 48°42’53”W; 6.7 ha). We ringed all territorial individuals in the study sites with color combinations to allow for individual identification. To capture individuals, we attracted them with playback and used 12-m-wide ornithological nets with 28-mm meshes.

**Figure 1.**
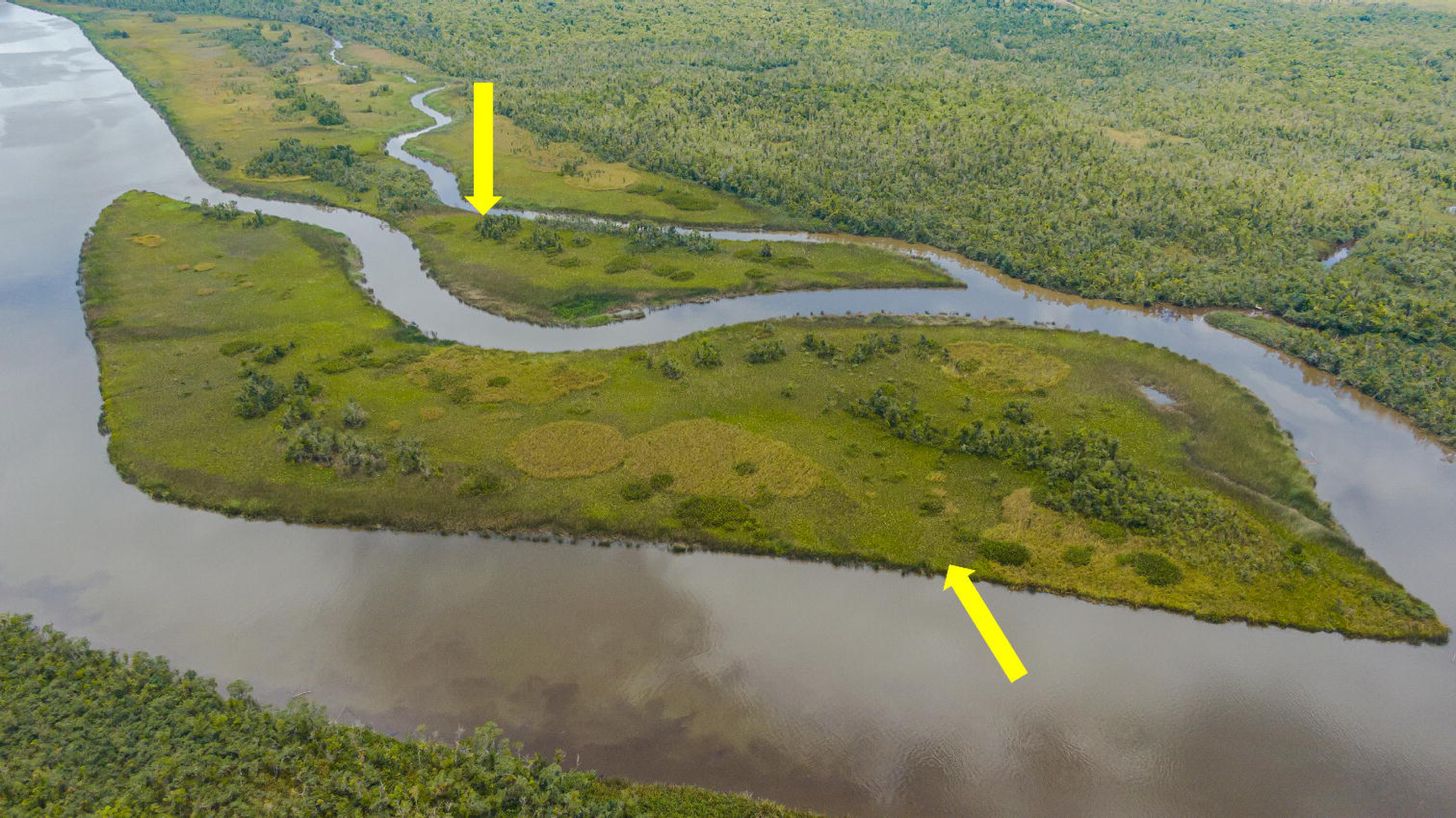
View of the study areas for *Formicivora acutirostris* in Continente (upper yellow arrow) and Jundiaquara Island (lower yellow arrow), Guaratuba Bay, Ramsar Site Guaratuba, in the Municipality of Guaratuba, Paraná coast, Southern Brazil. Photographed by Gabriel Marchi.

From January 2006 to mid-2008, we worked in the field every day from September to February, and six to eight days per month in the remaining months. From mid-2008, we worked in the field for three to eight days per month every month. Daily, we worked from dawn until about 12:00 or 1:00 pm and for an additional 2–3.5 h in the afternoon, before dusk. Fieldwork was conducted by three to six people.

We walked through all the study areas, marking the location of territorial individuals on each monthly fieldwork trip or every three days when the fieldwork was uninterrupted. Thus, we determined the number of territorial pairs in the study areas and assessed the densities of territorial individuals per 12-month cycle: 15 cycles on Jundiaquara Island (2006–2021), 14 cycles at Continente (2007–2021), 11 cycles on Folharada Island (2010–2021), and one cycle at Lagoa do Parado (2012–2013).

### Target environments

Jundiaquara Island, Folharada Island, and Continente are located in the Guaratuba Bay estuary, which is flooded twice daily floodings by mixed semi-diurnal tides (Lee and Chang 2019). In Lagoa do Parado, where the influence of tides is indirect, poor water drainage (Reinert *et al*. 2007) leads to periodic flooding during the rainy season (*c.* October to March). The studied habitats are summarized in Table S1.

### Estimation of the EOO

Since Reinert *et al*.’s (2007) study, we have found new sites of occurrence of the species in Paraná and Santa Catarina, where we conducted systematic searches. For each site, we used GPS to establish geographic coordinates and identified the species’ habitat, following Reinert *et al*. (2007). Additionally, we reviewed the species’ geographic distribution by compiling records from the literature and the Wiki Aves website (http://www.wikiaves.com.br). We joined the extreme points of occurrence to define the species’ EOO (*sensu* IUCN [2012]; see below).

### Estimation of the AOO

We analyzed the landscapes and mapped the habitat types used by the species (Reinert *et al*. 2007) within the EOO to determine the AOO. We mapped each of these habitats on 2020 orbital images from Google Earth Pro 7.3.3.7786 (64-bit), noting the positions, textures, height impressions, and color nuances of the vegetation. We checked *in loco* whether the mapping was correctly assigned to the classified habitats in the following regions: Antonina Bay (Antonina Municipality), Nhundiaquara River (Antonina and Morretes Municipalities), Guaratuba Bay (Guaratuba Municipality) in Paraná; Babitonga Bay (Garuva, Joinville, and São Francisco do Sul Municipalities), Itapoá, São Francisco do Sul, Laguna, and Jaguaruna Municipalities in Santa Catarina; and Dom Pedro de Alcântara Municipality in Rio Grande do Sul. There were situations in which we could not identify the dominant trees in habitats that had an upper arboreal stratum and a lower herbaceous stratum; hence, we treated them as “unidentified arboreal formations” (Table S1).

Based on the occupation area mapped by Reinert *et al*. (2007), we conducted evaluations for each habitat using Google Earth orbital images from 2002–2020 (Table 1), as follows: 1) whether the habitat still existed or had disappeared, and in the latter case, why it disappeared; 2) whether its area had changed; 3) whether the habitat had undergone progressive or regressive ecological succession, requiring reclassification as a different environment (*sensu* Sandretti-Silva *et al*. 2023); 4) whether the habitat had undergone progressive ecological succession to an environment not used by the species; and 5) whether the habitat had been misclassified. In the vicinity of the AOO mapped by Reinert *et al*. (2007), we also used orbital images from 2012 of Google Earth to conduct the following evaluations: 6) if new patches of habitats used by the species appeared, 7) whether an environment not formerly used by the species had undergone regressive succession to a used habitat, and 8) whether a habitat had been overlooked. We quantified the herbaceous formations suitable for the species but invaded by exotic grasses (*Urochloa arrecta* [*U. arrecta*] and *Urochloa mutica* [*U. mutica*]) as missing habitats (Reinert *et al*. 2007).

**Table 1.**
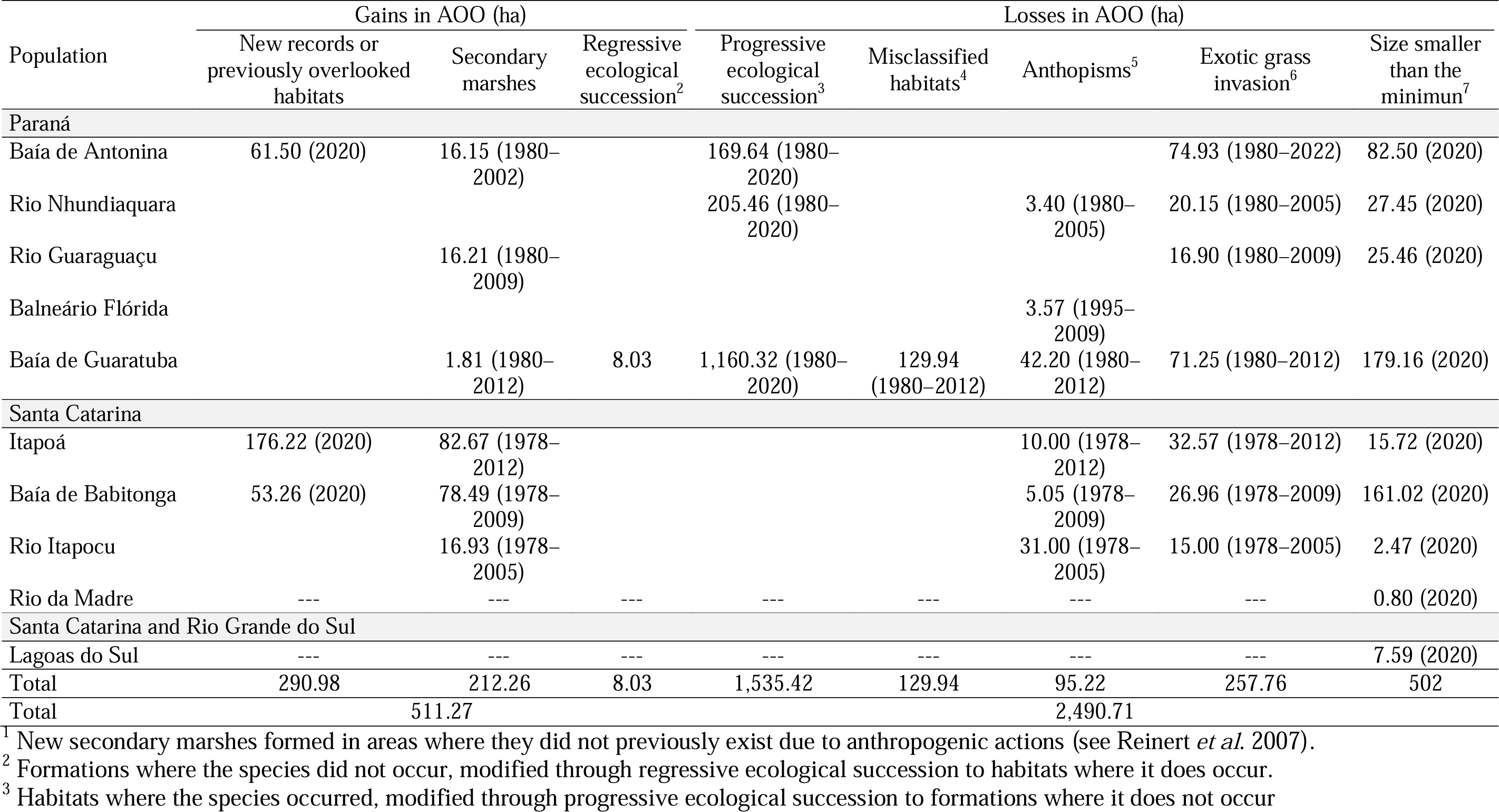
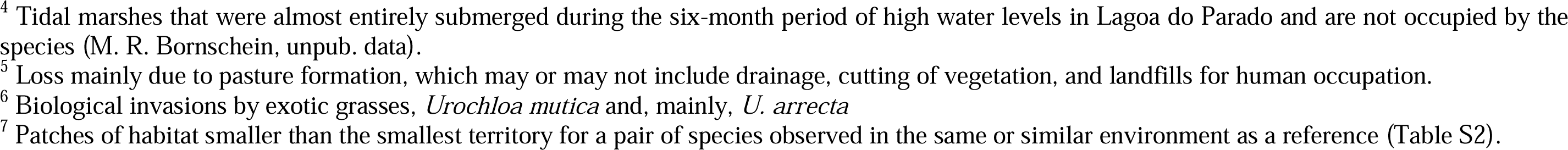
Gains and losses of the area of occupancy (AOO) by populations of the Paraná Antwren (*Formicivora acutirostris*) compared with the previous mapping (Reinert *et al*. 2007) based on aerial photographs from 1978 in Santa Catarina and from 1980 and 1995 in Paraná. The years or intervals of the years of the photographs or orbital images used for the analysis are indicated in parentheses.

| Population | Gains in AOO (ha) |  |  | Losses in AOO (ha) |  |  |  |  |
| --- | --- | --- | --- | --- | --- | --- | --- | --- |
|  | New records or previously overlooked habitats | Secondary marshes | Regressive ecological succession<sup>2</sup> | Progressive ecological succession<sup>3</sup> | Misclassified habitats<sup>4</sup> | Anthopisms<sup>5</sup> | Exotic grass invasion<sup>6</sup> | Size smaller than the minimum<sup>7</sup> |
| **Paraná** |  |  |  |  |  |  |  |  |
| Baía de Antonina | 61.50 (2020) | 16.15 (1980–2002) |  | 169.64 (1980–2020) |  |  | 74.93 (1980–2022) | 82.50 (2020) |
| Rio Nhundiaquara |  |  |  | 205.46 (1980–2020) |  | 3.40 (1980–2005) | 20.15 (1980–2005) | 27.45 (2020) |
| Rio Guaraguaçu |  | 16.21 (1980–2009) |  |  |  |  | 16.90 (1980–2009) | 25.46 (2020) |
| Balneário Flórida |  |  |  |  |  | 3.57 (1995–2009) |  |  |
| Baía de Guaratuba |  | 1.81 (1980–2012) | 8.03 | 1,160.32 (1980–2020) | 129.94 (1980–2012) | 42.20 (1980–2012) | 71.25 (1980–2012) | 179.16 (2020) |
| **Santa Catarina** |  |  |  |  |  |  |  |  |
| Itapoá | 176.22 (2020) | 82.67 (1978–2012) |  |  |  | 10.00 (1978–2012) | 32.57 (1978–2012) | 15.72 (2020) |
| Baía de Babitonga | 53.26 (2020) | 78.49 (1978–2009) |  |  |  | 5.05 (1978–2009) | 26.96 (1978–2009) | 161.02 (2020) |
| Rio Itapocu |  | 16.93 (1978–2005) |  |  |  | 31.00 (1978–2005) | 15.00 (1978–2005) | 2.47 (2020) |
| Rio da Madre | --- | --- | --- | --- | --- | --- | --- | 0.80 (2020) |
| **Santa Catarina and Rio Grande do Sul** |  |  |  |  |  |  |  |  |
| Lagoas do Sul | --- | --- | --- | --- | --- | --- | --- | 7.59 (2020) |
| Total | 290.98 | 212.26 | 8.03 | 1,535.42 | 129.94 | 95.22 | 257.76 | 502 |
| Total |  | 511.27 |  |  |  | 2,490.71 |  |  |

<sup>1</sup> New secondary marshes formed in areas where they did not previously exist due to anthropogenic actions (see Reinert *et al.* 2007).
<sup>2</sup> Formations where the species did not occur, modified through regressive ecological succession to habitats where it does occur.
<sup>3</sup> Habitats where the species occurred, modified through progressive ecological succession to formations where it does not occur
<sup>4</sup> Tidal marshes that were almost entirely submerged during the six-month period of high water levels in Lagoa do Parado and are not occupied by the species (M. R. Bornschein, unpub. data).
<sup>5</sup> Loss mainly due to pasture formation, which may or may not include drainage, cutting of vegetation, and landfills for human occupation.
<sup>6</sup> Biological invasions by exotic grasses, *Urochloa mutica* and, mainly, *U. arrecta*
<sup>7</sup> Patches of habitat smaller than the smallest territory for a pair of species observed in the same or similar environment as a reference (Table S2).

We were able to make comparisons with the areas of occupancy mapped by Reinert *et al*. (2007) using Google Earth orbital images from 2002–2020 because the previous mapping was printed on photocards (1:10,000) and digitized on Google Earth. This digitization was carried out by comparing the formations on the photocards with the older available Google Earth satellite images.

We treated the new mapping with two peculiarity. The first peculiarity was on the central and southern coast of Santa Catarina and on the north coast of Rio Grande do Sul, specifically south of 27°45’S to the southernmost record of the species, where we mapped only the habitats with an upper arboreal stratum and a lower herbaceous stratum where we observed the species’ occurrence, leaving out the herbaceous formations: tidal marshes (*brejo de maré*), saw grass marshes (*brejo de capim-serra*), meander marshes (*brejo de meandro*), marshes between coastal dunes (*brejo intercordão*), and secondary marshes (according to Reinert *et al*. [2007]). The second peculiarity was to remove from the mapping of the AOO the patches of habitat smaller than the smallest territory for a species pair observed in the same or in a similar reference habitat (Table S2). We evaluated a set of patches and summed their areas if they were up to 6 m apart. In the case of different habitat patches, they were considered if the total sum of their area ratios compared to the reference territory’s areas resulted in a value equal to or greater than 1. We used this distance because 6 m is the largest gap of other habitats that we recorded in a species territory (M. R. Bornschein, unpub. data).

We summed all the areas of the mapped habitats by habitat type, following Reinert *et al*. (2007). The total was the AOO (IUCN 2012).

### Delimitation of distinct populations

New sites of records were either assigned to one of the existing eight populations of the species or assigned to a new population if the habitat patches were close to each other and at least 10 km away from another set of habitat patches (Reinert *et al*. 2007). Within this distance of at least 10 km between patches, we confirmed the absence or inferred absence of patches in highly degraded regions where natural habitats have been suppressed. We then considered the existence of estuaries, meandering rivers flowing into estuaries or lagoons, and lagoons in plains bordered by vast flat areas as indicating the possible past presence of habitats used by the species.

### Estimation of population size

To estimate the population size of *F. acutirostris*, we multiplied the values of the density of territorial individuals (Table 2) by the total area of the same habitat, following Reinert *et al*. (2007). We obtained territory size data from the literature (Reinert *et al*. 2007) and transformed them into densities based on the size of the studied area and the number of existing territories (Table 2). Some sites were studied for several years, so we averaged the densities of individuals (Tables S3–S5). For habitats where density data were not available, we applied densities obtained in habitats with similar floristic structures (Table 3; adapted from Reinert et al. [2007]).

**Table 2.**
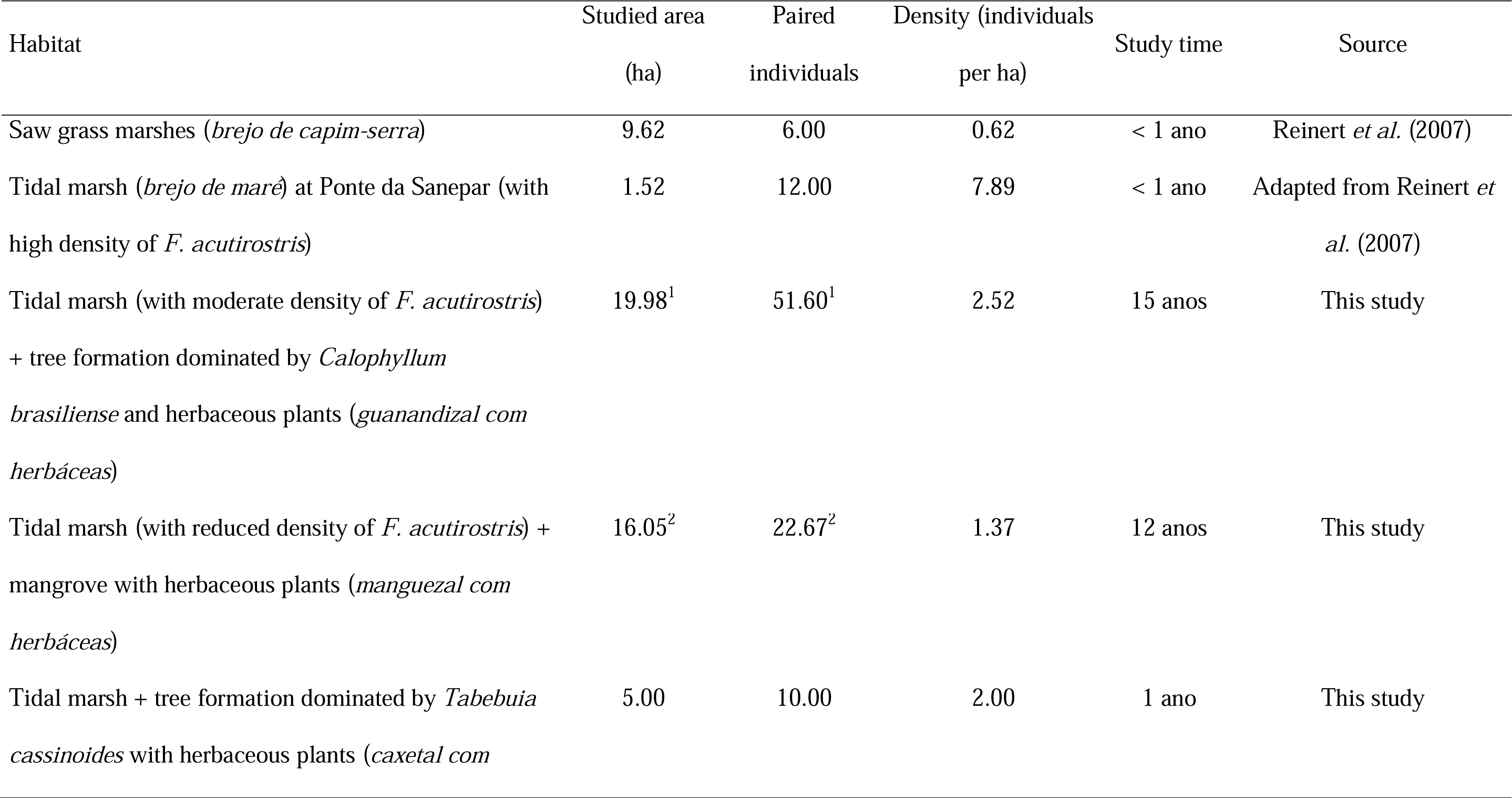
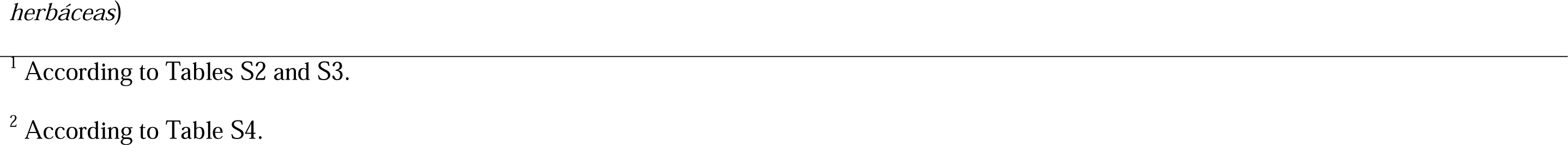
Average densities of territorial individuals of *Formicivora acutirostris* in the studied habitats in Guaratuba Bay, Paraná State, Southern Brazil, used for population size calculation.

| Habitat | Studied area<br>(ha) | Paired<br>individuals | Density (individuals<br>per ha) | Study time | Source |
| --- | --- | --- | --- | --- | --- |
| Saw grass marshes (*brejo de capim-serra*) | 9.62 | 6.00 | 0.62 | < 1 ano | Reinert *et al.* (2007) |
| Tidal marsh (*brejo de maré*) at Ponte da Sanepar (with high density of *F. acutirostris*) | 1.52 | 12.00 | 7.89 | < 1 ano | Adapted from Reinert *et al.* (2007) |
| Tidal marsh (with moderate density of *F. acutirostris*) + tree formation dominated by *Calophyllum brasiliense* and herbaceous plants (*guanandizal com herbáceas*) | 19.98<sup>1</sup> | 51.60<sup>1</sup> | 2.52 | 15 anos | This study |
| Tidal marsh (with reduced density of *F. acutirostris*) + mangrove with herbaceous plants (*manguezal com herbáceas*) | 16.05<sup>2</sup> | 22.67<sup>2</sup> | 1.37 | 12 anos | This study |
| Tidal marsh + tree formation dominated by *Tabebuia cassinoides* with herbaceous plants (*caxetal com* | 5.00 | 10.00 | 2.00 | 1 ano | This study |

<sup>1</sup> According to Tables S2 and S3.
<sup>2</sup> According to Table S4.

**Table 3.** Estimate of area of occupancy (AOO; ha) and population size of mature territorial individuals of *Formicivora acutirostris* by habitat type.

| Habitat | Estimated or<br>applied<br>density<sup>1</sup> | Populations |  |  |  |  |  |  |  |  |  | Total |
| --- | --- | --- | --- | --- | --- | --- | --- | --- | --- | --- | --- | --- |
|  |  | Paraná (PR) |  |  |  |  | Santa Catarina (SC) |  |  |  | SC and RS |  |
|  |  | Baía de | Rio | Rio | Balneário | Baía de | Itapoá | Baía de | Rio | Rio da | Lagoas do |  |
|  |  | Antonina | Nhundiaquara | Guaraguaçu | Flórida | Guaratuba |  | Babitonga | Itapocu | Madre | Sul |  |
| Saw grass marshes (*brejo de capim-serra*) | 0.62 |  |  | 38.28 |  | 30.37 | 18.98 | 80.79 |  |  |  | 168.42 |
| Tidal marsh (*brejo de maré*) at Ponte da | 7.89 |  |  |  |  | 4.82 |  |  |  |  |  | 4.82 |
| Sanepar (with high density of *F. acutirostris*) |  |  |  |  |  |  |  |  |  |  |  |  |
| Tidal marsh (with high or moderate density of *F. acutirostris*) | 2.52 | 37.24 | 0.80 | 7.95 |  | 101.08 | 1.13 | 209.77 | 5.17 |  |  | 363.14 |
| Tidal marsh (with low density of *F. acutirostris*) | 1.37 | 51.53 | 86.85 |  |  | 14.23 |  | 12.03 |  |  |  | 164.64 |
| Marsh between coastal dunes (*brejo intercordão*) | 2.52 |  |  |  | 1.97 |  | 4.29 |  |  |  |  | 6.26 |
| Meander marsh (*brejo de meandro*) | 2.52 | 1.60 |  |  |  |  |  |  | 22.40 |  |  | 24.00 |
| Secondary marsh | 2.52 | 221.6 |  |  |  | 52.39 | 70.50 | 43.73 | 29.36 |  | 7.11 | 424.69 |
| Tidal marsh + tree formation dominated by *Tabebuia cassinoides* with herbaceous plants (*caxetal com herbáceas*) | 2.00 |  |  | 4.70 |  | 472.35 |  |  |  |  |  | 477.05 |
| Tidal marsh + tree formation dominated by *Calophyllum brasiliense* with herbaceous plants (*guanandizal com herbáceas*) | 2.52 | 29.89 | 42.37 |  |  | 87.48 |  |  |  |  |  | 159.74 |
|  |  | Paraná (PR) |  |  |  |  | Santa Catarina (SC) |  |  |  | SC and RS |  |
|  |  | Baía de | Rio | Rio | Balneário | Baía de | Itapoá | Baía de | Rio | Rio da | Lagoas do |  |
|  |  | Antonina | Nhundiaquara | Guaraguaçu | Flórida | Guaratuba |  | Babitonga | Itapocu | Madre- | Sul |  |
| Tidal marsh + mangrove with herbaceous plants (*manguezal com herbáceas*) | 1.37 | 121.56 | 31.79 | 3.98 |  | 93.60 |  | 512.39 | 13.61 |  |  | 776.93 |
| Tidal marsh + tree formation dominated by *C. brasiliense* with herbaceous plants + mangrove with herbaceous plants | 2.52 | 104.40 |  |  |  | 36.35 |  |  |  |  |  | 140.75 |
| Tidal marsh + unidentified arboreal formation | 2.00 |  |  |  |  |  | 51.00 |  |  |  |  | 51.00 |
| Secondary marsh + unidentified arboreal formation | 2.00 | 22.60 |  |  |  |  |  |  |  |  |  | 22.60 |
| Tree formation dominated by *T. cassinoides* with herbaceous plants | 0.62 | 103.87 |  | 10.00 |  | 34.71 |  |  |  |  |  | 148.58 |
| Tree formation dominated by *C. brasiliense* with herbaceous plants | 0.62 | 45.53 | 78.20 | 22.70 |  | 184.60 |  |  |  |  |  | 285.5 |
| Tree formation dominated by *C. brasiliense* with herbaceous plants (regressive ecological succession) | 0.62 |  |  |  |  | 8.03 |  |  |  |  |  | 53.56 |
| Mangrove with herbaceous plants | 0.62 | 285.60 | 51.35 | 27.92 |  | 90.67 | 4.48 | 61.17 |  |  |  | 521.19 |
| Tree formation dominated by *C. brasiliense* with herbaceous plants + mangrove with | 0.62 | 13.20 |  |  |  | 56.48 |  |  |  |  |  | 69.68 |
|  |  | Paraná (PR) |  |  |  |  | Santa Catarina (SC) |  |  |  | SC and RS |  |
|  |  | Baía de | Rio | Rio | Balneário | Baía de | Itapoá | Baía de | Rio | Rio da | Lagoas do |  |
|  |  | Antonina | Nhundiaquara | Guaraguaçu | Flórida | Guaratuba |  | Babitonga | Itapocu | Madre | Sul |  |
| herbaceous plants |  |  |  |  |  |  |  |  |  |  |  |  |
| Unidentified arboreal formation | 0.62 |  |  |  |  |  | 13.12 |  |  |  |  | 13.12 |
| Unidentified arboreal formation (south of<br>27°45'S)<sup>2</sup> | 1.37 |  |  |  |  |  |  |  |  | 2.06 | 151.5 | 153.56 |
| Unidentified arboreal formation in succession<br>of secondary marsh | 0.62 | 72.66 |  |  |  |  |  |  |  |  |  | 72.66 |
| Total area of occupancy |  | 1,111.28 | 291.36 | 115.53 | 1.97 | 1,267.16 | 163.50 | 919.88 | 70.54 | 2.06 | 158.61 | 4,101.89 |
| Total number of individuals |  | 1,599.99 | 351.65 | 96.20 | 4.96 | 2,080.27 | 316.00 | 1,445.29 | 162.11 | 2.82 | 225.47 | 6,284.76 |

<sup>1</sup> According to Table 2.
<sup>2</sup> Field data for the Rio da Madre and Lagoas do Sul populations suggest a higher density than 0.62 individuals per hectare, which is the density value for similar habitats in other populations.

### Conservation Status and Green Status

We evaluated the conservation status of the species according to IUCN (2012, 2022) and the Green Status according to the IUCN Species Conservation Success Task Force (2020) and IUCN (2021). We obtained the Current, Counterfactual Current, Future-with-conservation, and Future-without-conservation scenarios to assess the metrics for Conservation Legacy, Conservation Dependence, and Conservation Gain (IUCN 2021). We developed future scenarios by extrapolating the annual decline rates for habitats and individuals from populations and assessing threats in the areas.

## Results

### AOO

The estimated AOO of *F. acutirostris* was 4,101.89 ha (Table 3), distributed across 10 populations (Figure 2A), including two new populations named Rio da Madre and Lagoas do Sul (Table 2). We recorded the species at new sites, such as the Faisqueira River (25°21’58”S, 48°38’42”W), which is part of the Baía de Antonina population in Paraná, and the other three areas hosted by the Itapoá population in Santa Catarina (26°07’53”S, 48°39’26”W; 26°10’51”S, 48°36’53”W; 26°09”°23”S, 48°37’50”W). The Baía de Babitonga population in Santa Catarina had also expanded due to the detection of habitat types used by the species that were overlooked in the previous mapping (Table 1). In the new population Lagoas do Sul, we recorded the species in Pedras Brancas (28°30’54”S, 48°47’29”W), municipality of Laguna, in the state of Santa Catarina; and Lagoa do Forno (29°22’56”S, 49°53’12”W), municipality of Dom Pedro de Alcântara, in the state of Rio Grande do Sul.

**Figure 2.**
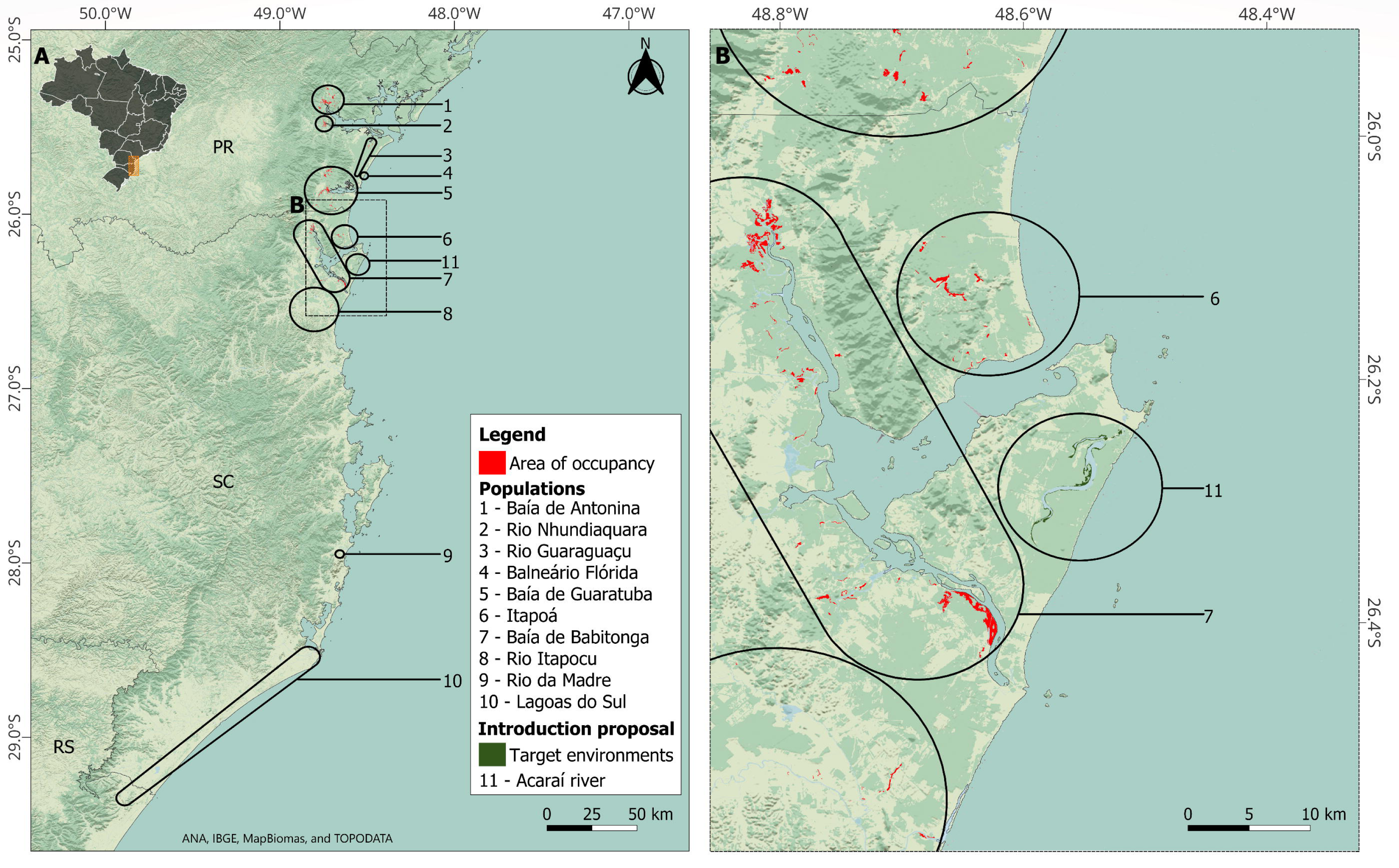
A. Area of occupancy of *Formicivora acutirostris* in its 10 populations. B. We highlighted the target areas for the assisted colonization of the species along the Acaraí River, on the northern coast of Santa Catarina (Population 11). Abbreviations: PR = Paraná; SC = Santa Catarina; RS = Rio Grande do Sul. Background images from National Water and Sanitation Agency (ANA), Brazilian Institute of Geography and Statistics (IBGE), MapBiomas, and TOPODATA.

We identified a 146-km distributional gap between the southernmost previously mapped population (Rio Itapocu) and the northernmost of the two new populations (Rio da Madre). We searched unsuccessfully for the species in various areas for different years within this gap and within the Rio da Madre and Lagoas do Sul populations (1998–2012), but in herbaceous formations dominated by the bulrush *Schoenoplectus californicus*, swamp saw grass *Cladium mariscus*, southern cattail *Typha domingensis*, or *Scirpus giganteus*. These surveys were carried out in several municipalities, such as Araranguá, Itajaí, Garopaba, Jaguaruna, Laguna, Navegantes, Palhoça, Paulo Lopes, and Penha in Santa Catarina, and Dom Pedro de Alcântara in Rio Grande do Sul. However, following the first records of *F. acutirostris* in the Laguna Municipality in early 2012, subsequent records by the authors, other researchers, and bird observers have indicated that the species’ habitat in the two southern populations is dominated by the mangrove fern *Acrostichum danaeifolium,* together with the trees Brazilian pepper tree *Schinus terebinthifolius* and *Myrsine parvifolia*. For example, all the photographs of the species on Wiki Aves (74 photographs up to 30 May 2023) obtained from the Rio da Madre (Paulo Lopes municipality) and Lagoas do Sul (Laguna and Jaguaruna municipalities) populations revealed the species in a habitat exclusively dominated by *A. danaeifolium* or in association with *S. terebinthifolius* and/or *M. parvifolia*. However, our observations at Lagoa do Forno were in a flooded formation dominated by *T. domingensis* or by this plant alongside trees. The 16 photographs of *F. acutirostris* in Rio Grande do Sul on the Wiki Aves platform (up to 30 May 2023), probably taken at Lagoa do Forno, show the species in a habitat characterized by the presence of the herbaceous plants *T. domingensis*, *S. californicus*, *Fuirena* sp., and at least some Melastomataceae shrubs. The numerous anthropogenic impacts on Lagoa do Forno, including landfills for roads, drainage canals, and deforestation of tree vegetation, indicate that the habitat has become degraded, with a greater abundance of herbaceous plants and a diminished presence of trees, and that the presence of *F. acutirostris* in this habitat may be a local adaptation or colonization in response to these changes.

A considerable part of the AOO of *F. acutirostris* is susceptible to tidal flooding (92.7% - excluding marginal habitats in secondary marshes); the remaining portion (including secondary marshes) is located at low altitudes near rivers or in lagoons. We regard the species as endemic to salt marshes, occurring across a broad range of floristic composition along the gradient of altitude and salinity in tide-influenced areas. These habitats range from highly saline habitats with some herbaceous species to low- or no-salinity habitats with a lower stratum characterized by typical marsh herbaceous plants and an upper stratum containing trees commonly found in flood-prone areas (see Table S1). One notable aspect is that the eight populations of the species in Paraná and Northern Santa Catarina occurred across the entire floristic gradient, whereas the two southern populations occurred only at the end of the gradient in formations with an upper arboreal stratum of tree species common to flood-prone areas and a lower herbaceous stratum with typical marsh plants, although some observations in Rio Grande do Sul deviated from this pattern, possibly indicating adaptation to local anthropogenic impacts (see previous paragraph).

Changes occurred in the species’ AOO between 1980 and 2020 due to nine factors, including gains according to new records and losses from invasion by exotic grasses (*U. arrecta* and *U. mutica*; Table 1). Throughout this period, there were gains in the AOO due to the formation of new habitats, but their sizes were still smaller than the minimum size for a territory in a similar habitat (tidal marsh with reduced species density; Table S2), so this change is not accounted for in Table 1. Another gain in AOO occurred due to regressive ecological succession, with tree mortality and the establishment of herbaceous species in the lower stratum (Table 1). We detected an area of 22.17 ha with this phenomenon in the vicinity of Baía de Guaratuba population (Table 1), always in elevated positions within the tidal plain between herbaceous-dominated formations (saw grass marshes) and forests. However, only 8.03 ha of these habitats formed through regressive succession met the minimum size of territory for the species (Tables 1 and S2). This phenomenon may be attributed to higher tides increasing salinity in those areas.

There was a small loss of AOO due to human activities between 1980 and 2020 (95.22 ha; Table 1), contrary to our initial expectations. However, this assessment did not consider the Rio da Madre and Lagoas do Sul populations (Table 1), which could have experienced significant losses, but the impacts in these regions are severe and longstanding and require analysis of the landscape based on aerial or orbital images from before 1980. The most notable loss of AOO of *F. acutirostris* occurred through progressive ecological succession of habitats that had both arboreal and herbaceous stratum with typical marsh plants in 1980 and succeeded to environments without the typical marsh plants in the lower stratum and that were no longer occupied by the species by 2020 (1,535.42 ha; Table 1). This phenomenon is natural and expected. The most significant impact we detected on populations was the loss of AOO due to invasion by the exotic grasses *U. arrecta* and *U. mutica*. In the field, we observed the presence of one or both species associated in all populations, except in Rio da Madre. Areas where exotic species are present but at low densities cannot be distinguished from aerial or orbital images, whereas areas dominated by exotic species can be easily identified. We observed areas where that grasses were dominant in association with seven populations (Table 1). The loss of habitat due to this factor was not accounted for the Balneário Flórida population as the invaded habitats were filled in 2009, so the impact was considered a result of human activities (Table 1).

The exotic grasses started to spread over the AOO of *F. acutirostris* from rural properties upstream, where riparian vegetation had been eliminated. *Urochloa arrecta* forms large masses that spread as floating vegetation on rivers, sometimes accompanied by smaller amounts of *U. mutica*. These floating masses are occasionally detached by river floods and carried to other areas, eventually reaching downstream habitats of *F*. *acutirostris*. This spread occurred after 1980, since no AOO of *F. acutirostris* was impacted by the dominance of exotics before that date. Another cause of the spread of exotic grasses is human-mediated transportation, both indirect, though boats and fishing gear (e.g., nets and tanks with live bait), and direct, through the planting of exotic grasses along riverbanks to attract capybaras *Hydrochoerus hydrochaeris* for hunting with traps. We did not detect vegetative propagation of the exotic grasses, at least in the estuaries, where seed banks do not form, possibly due to salinity limitations (observation made since the beginning of the management of these exotic species in the estuary of Guaratuba Bay in 2012).

The floating masses of *U. arrecta* carried by ebb tides became established in the habitats of *F. acutirostris* with superficial water salinity levels up to 11 ‰ (e.g., 25°51’56”S, 48°32’03”W; unpublished data collected by the authors in 2012). However, after an extreme weather event in the winter of 2020 (a bomb cyclone), which may have caused a greater influx of saltwater into the estuary, several patches dominated by this exotic species downstream of the estuary died and did not regenerate. We observed this mortality in patches of *U. arrecta* in the vicinity of Baía de Antonina (through analysis of orbital images) and Baía de Guaratuba (*in loco*) populations.

We detected six regions that had large, concentrated patches of *F. acutirostris* minimally invaded by *U. arrecta and U. mutica* (Table 4, Figure 3), associated with four populations of the species. These regions account for a total of 1,945.38 ha (47.4% of the total AOO of the species; Table 4).

**Table 4.** The smallest area less impacted by the invasion of the exotic grasses *Urochloa arrecta* and *U. mutica*. They are proposed here as strategic units for management and conservation, called exotic-free zones (EFZs).

| EFZ | Species population | Estimated area (ha) | Estimated population (mature territorial individuals) |
| --- | --- | --- | --- |
| Confluence of Cacatu and Lagoa Vermelha Rivers | Baía de Antonina | 444.04 | 601.17 |
| Confluence of Nhundiaquara and Neves Rivers | Rio Nhundiaquara | 161.67 | 206.61 |
| Lagoa do Parado | Baía de Guaratuba | 228.56 | 457.12 |
| Confluence of São João and Cubatão Rivers | Baía de Guaratuba | 431.46 | 643.25 |
| Palmital River | Baía de Babitonga | 335.27 | 512.76 |
| da Passagem Island | Baía de Babitonga | 344.38 | 631.24 |
| Total |  | 1,945.38 | 3,052.15 |

**Figure 3.**
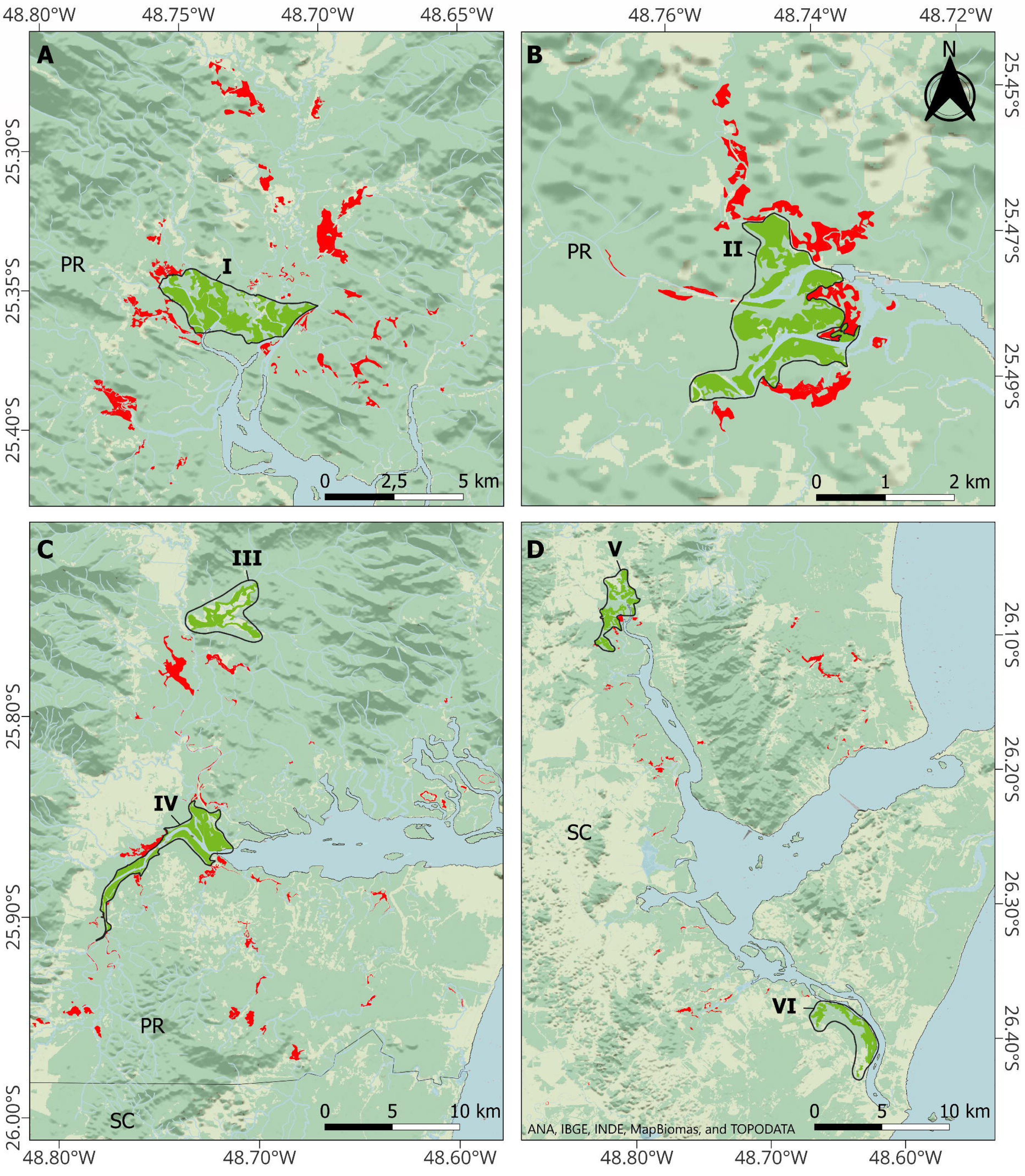
The smallest area of area of occupancy (in red and green) containing a significant range of environments that were less impacted by the invasion of the exotic grasses *Urochloa arrecta* and *U. mutica*. We propose that these areas be considered *exotic-free zones* (EFZs)—a concept for landscape management that offers advantageous costs versus benefits. A. I. EFZ Confluence of the Cacatu and Lagoa Vermelha Rivers (Baía de Antonina population). B. II. EFZ Confluence of the Nhundiaquara and Neves Rivers (Rio Nhundiaquara population). C. III. EFZ Lagoa do Parado, (Baía de Guaratuba population). C. IV. EFZ Confluence of the São João and Cubatão Rivers (Baía de Guaratuba population). D. V. Palmital River (Baía de Babitonga population). D. VI. da Passagem Island (Baía de Babitonga population). Abbreviations: PR = Paraná; SC = Santa Catarina. Background images from ANA, IBGE, INDE, MapBiomas, and TOPODATA.

### Estimation of population size

We estimated the population size of the species to be 6,284 individuals (Table 3), 4,133 of which were present in Paraná. We estimated the largest population to have 2,080 individuals (Baía de Guaratuba population, in Paraná), and the smallest population to have two individuals (Rio da Madre population, in Santa Catarina; Table 3). The population estimates for the six regions with large, concentrated patches of minimally invaded habitats are presented in Table 4.

### Conservation status

The estimation of how the losses and gains of habitat (Table 1) affected the losses and gains of individuals indicated an annual loss of 44.76 individuals (Table 5), totaling a loss of 644.54 individuals over three generations (10.31% of the total population, with each generation lasting 4.8 years; BirdLife International [2019]). Since we estimated the population to be less than 10,000 mature individuals (Table 3) and considering the estimated loss of 10% over three generations, the species should be classified as Vulnerable (VU C1).

**Table 5.**
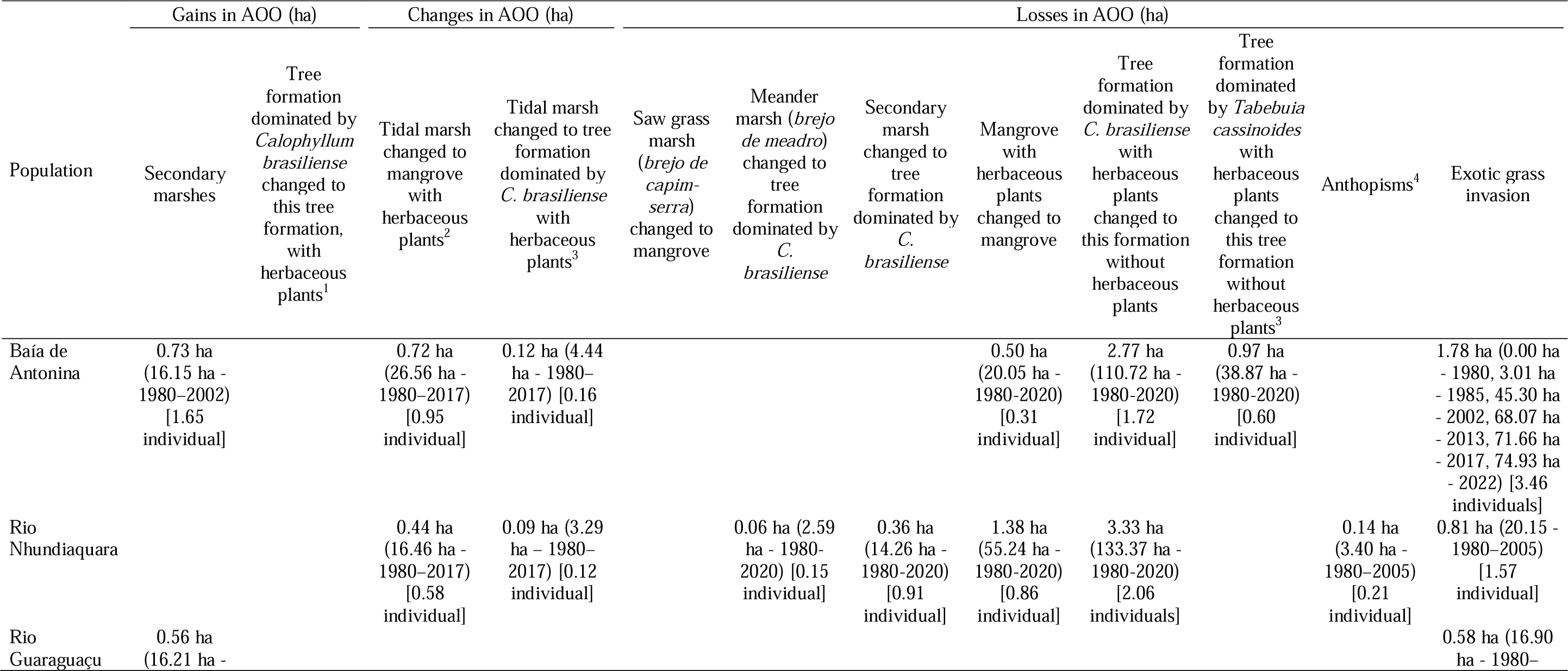
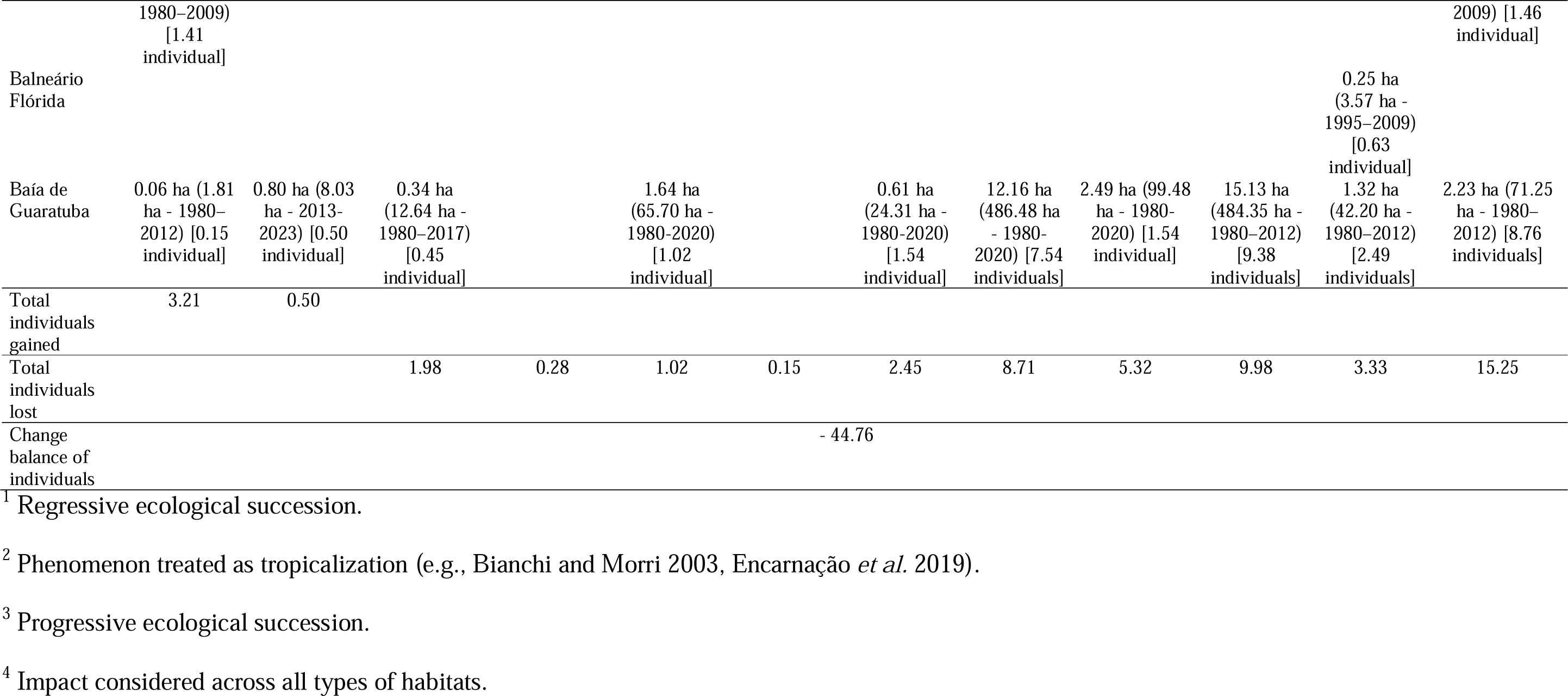
Estimated gains or losses of individuals of *Formicivora acutirostris* due to changes in the area of occupancy (AOO) of the populations where both values could be calculated. The following values are presented: average annual change in area of occupancy; total changed area, intervals of years of this evaluation (in parentheses); and estimated number of mature territorial individuals gained or lost annually (in brackets). To calculate the impact on the individuals, we multiplied the average annual change in area by the average density of individuals in the respective habitats, as shown in Table 4. When the changes in area of occupancy involved ecological succession from one habitat to another, we multiplied the average annual change in area by the obtained value for the decreased density of individuals of *F. acutirostris* in each of these habitats (according to Table 3).

### Green Status

The Current scenario showed that *F. acutirostris* is Largely Depleted, with a Species Recovery Score of 42.5%. In this scenario, only three of the 10 populations were not facing a specific threat and could be deemed Functional despite their naturally small population sizes. Of the remaining populations, three larger populations were assessed as Endangered (EN) in terms of area or number of locations, and four as Critically Endangered (CR; Table 6).

**Table 6.**
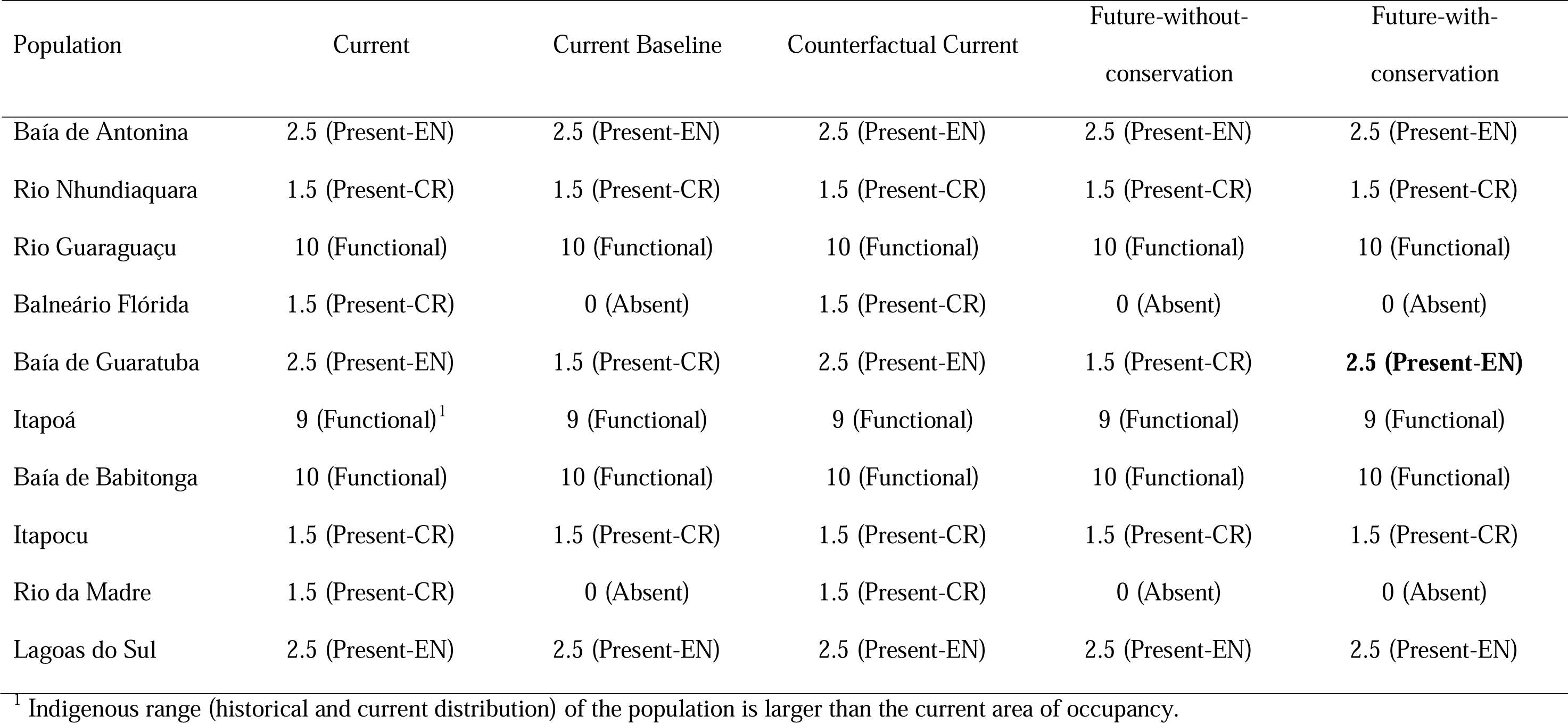
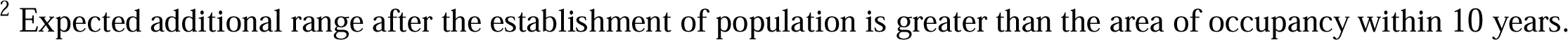
Evaluation of the Green Status of *Formicivora acutirostris* across its 10 populations and an eleventh (Acaraí river), if the suggested assisted colonization occurs at this site, according to IUCN Species Conservation Success Task Force (2020) and IUCN (2021). The fine-resolution weights and categories for the spatial units and the resulting Green Score for the scenarios are presented. Bold values indicate the values that have changed compared to the Current Baseline scenario.

| Population | Current | Current Baseline | Counterfactual Current | Future-without-conservation | Future-with-conservation |
| --- | --- | --- | --- | --- | --- |
| Baía de Antonina | 2.5 (Present-EN) | 2.5 (Present-EN) | 2.5 (Present-EN) | 2.5 (Present-EN) | 2.5 (Present-EN) |
| Rio Nhundiaquara | 1.5 (Present-CR) | 1.5 (Present-CR) | 1.5 (Present-CR) | 1.5 (Present-CR) | 1.5 (Present-CR) |
| Rio Guaraguaçu | 10 (Functional) | 10 (Functional) | 10 (Functional) | 10 (Functional) | 10 (Functional) |
| Balneário Flórida | 1.5 (Present-CR) | 0 (Absent) | 1.5 (Present-CR) | 0 (Absent) | 0 (Absent) |
| Baía de Guaratuba | 2.5 (Present-EN) | 1.5 (Present-CR) | 2.5 (Present-EN) | 1.5 (Present-CR) | **2.5 (Present-EN)** |
| Itapoá | 9 (Functional)<sup>1</sup> | 9 (Functional) | 9 (Functional) | 9 (Functional) | 9 (Functional) |
| Baía de Babitonga | 10 (Functional) | 10 (Functional) | 10 (Functional) | 10 (Functional) | 10 (Functional) |
| Itapocu | 1.5 (Present-CR) | 1.5 (Present-CR) | 1.5 (Present-CR) | 1.5 (Present-CR) | 1.5 (Present-CR) |
| Rio da Madre | 1.5 (Present-CR) | 0 (Absent) | 1.5 (Present-CR) | 0 (Absent) | 0 (Absent) |
| Lagoas do Sul | 2.5 (Present-EN) | 2.5 (Present-EN) | 2.5 (Present-EN) | 2.5 (Present-EN) | 2.5 (Present-EN) |

<sup>1</sup> Indigenous range (historical and current distribution) of the population is larger than the current area of occupancy.
<sup>2</sup> Expected additional range after the establishment of population is greater than the area of occupancy within 10 years.

The Counterfactual Current scenario was similar to the Current scenario for all spatial units (Table 6). Therefore, the Conservation Legacy of the actions implemented for the species thus far is Zero (0%). Among the considered actions were the creation of integral protection conservation units (Bom Jesus Biological Reserve, for the Baía de Antonina population; Guaraguaçu Ecological Station, for the Guaraguaçu population; and the State Park Boguaçu, for the Baía de Guaratuba population), together with a project to translocate eggs between the Baía de Guaratuba and Rio Nhundiaquara populations and programs to eradicate the exotic grasses in the Baía de Guaratuba population (2012–2013 and 2012–2021; Figure 4).

**Figure 4.**
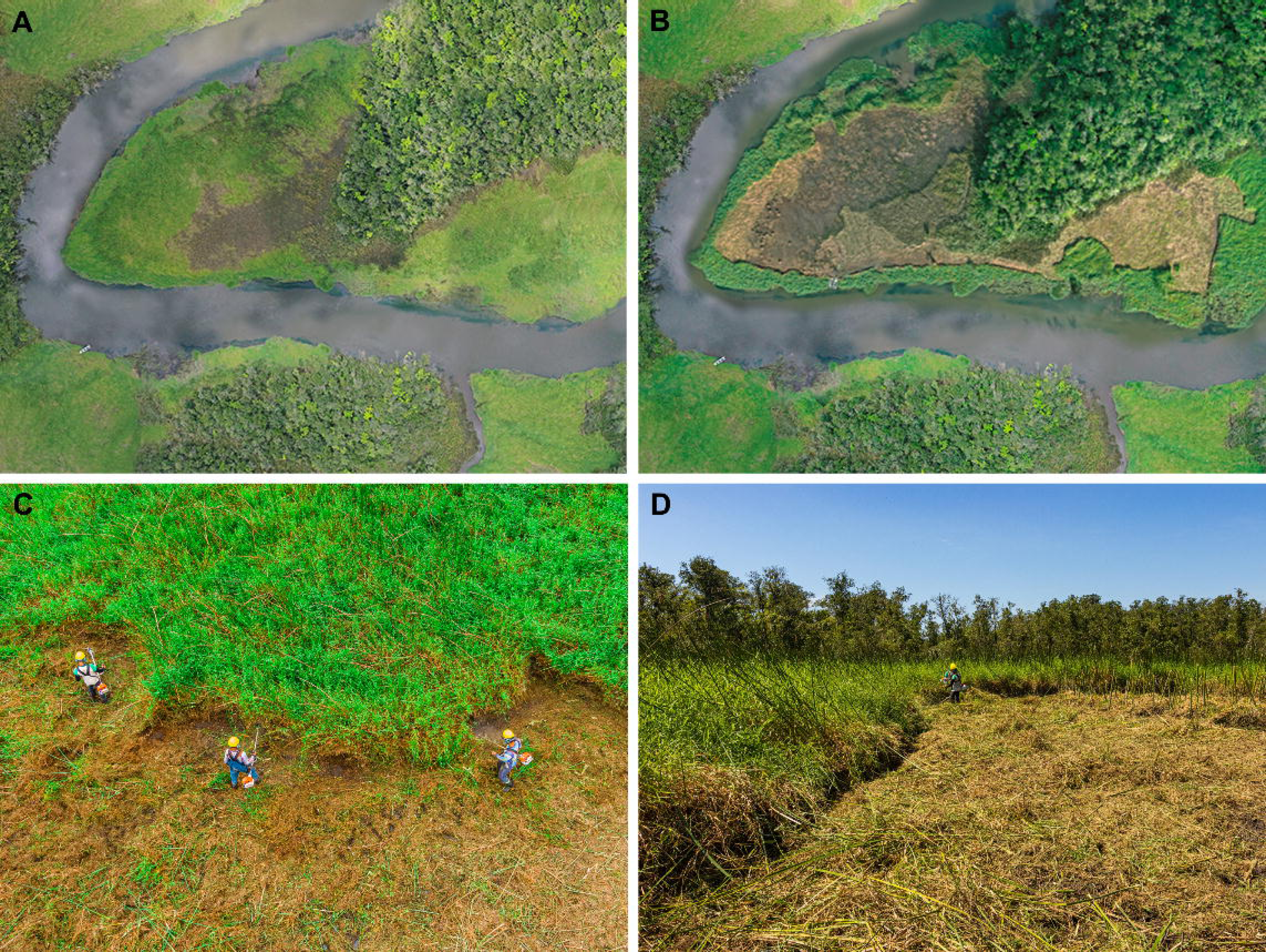
*Formicivora acutirostris* areas under management to eradicate the exotic grass *Urochloa arrecta*. A. Area invaded by exotic grasses prior to management. B. The central region of the same area with management, with biomass piled and stacked to prevent it from being carried by high tides. C. Management by clear cutting of vegetation. D. Cut biomass prior to piling and stacking. Photographed by Larissa Teixeira (A and B) and Gabriel Marchi (C and D).

In the Current Baseline and Future-without-conservation scenarios the species was assessed as Largely Depleted, with a Green Score of 38.5%. Therefore, the Conservation Dependency of the ongoing actions is Zero (0%). Currently, an exotic grass eradication program is being implemented in the Baía de Guaratuba population, but it is unable to improve the Green Score for the Current Baseline scenario due to its small scale (6 ha). In both scenarios, the Balneário Flórida and Rio da Madre populations became extinct because individual losses over 10 years exceeded the current population size (Table 7). Additionally, the Baía de Guaratuba population, which hosts the largest number of individuals of the species (Table 3), could be classified as Critically Endangered (CR) due to a decline in habitat, estimated at 350 ha in 10 years (based on Table 5).

In the future-with-conservation scenario the species was also assessed as Largely Depleted, with a Green Score of 43.2%. Consequently, the Conservation Gain via the planned conservation actions is Low (4.7%), but High considering substantial recovery (12.2% of the Green Score of the Current Baseline scenario) with the potential to increase in a few years. In this scenario, the Balneário Flórida and Rio da Madre populations continued to face extinction without planned conservation action to maintain individuals in such small and degraded areas (see Table 3). Future management actions may achieve satisfactory conservation results. The eradication of exotic grasses in the Baía de Guaratuba population at a rate of 8.5 ha per year could prevent its assessment as Critically Endangered. In addition, assisted colonization would add a functional spatial unit to 40% of its area (Table 6), but with the potential to achieve full functionality following recruitment of juveniles in successive years, indicating the establishment of the population (Figure 2B).

## Discussion

### Geographic distribution

Updating the geographic distribution, AOO, and population size of *F. acutirostris* was extremely timely (Reinert and Bornschein 2008, Reinert *et al*. 2009) because it occurs in coastal areas of Brazil experiencing intense urban expansion (e.g., cities of Paranaguá, Guaratuba, and Joinville), and because most of Reinert *et al*.’s (2007) mapping was based on aerial photographs taken over 40 years ago. Reinert *et al*. (2007) raised the possibility that the species could be recorded further south of the previously known southern limit, but its occurrence as far as 329 km south was unexpected (M. R. Bornschein, unpub. data). However, it is possible that the species occurs even further south than the currently mapped southern limit (29°23’04”S, 49°53’10”W, municipality of Dom Pedro de Alcântara in Rio Grande do Sul); for example, in the margins of lagoons with herbaceous and arboreal vegetation in the municipality of Osório in Rio Grande do Sul (29°51’43”S, 50°13’19”W). These areas were not considered part of the AOO of *F. acutirostris* because they were outside the extent of occurrence (see the Methods section).

### AOO and population size

The new AOO of the species is estimated to be 32.3% smaller than the previous estimate (6,060 ha; Reinert *et al*. [2007]), despite the discovery of two new populations, and the new estimate of population size is 64.5% smaller than the previous estimate (17,680 mature individuals; Reinert *et al*. [2007]), mainly due to two factors that led to overestimation of the population in the previous study. In the present study, we used the density of individuals in the study areas rather than the average territory area to calculate the population estimate, which incorporated “leftover” areas between territories that were perhaps too small or low quality to support a species pair. Reinert *et al*. (2007) considered every area of habitat to be occupied by the species. The second factor was the new density in other habitats (Table 2), revealing that the density of 7.89 individuals per hectare (Reinert *et al*. 2007) was high and restricted to a small area in the Baía de Guaratuba population (in tidal marshes with high floristic richness and reduced tidal flooding).

### Losses of AOO

The habitats of *F. acutirostris*, both in herbaceous formations and formations with an upper arboreal stratum and a lower herbaceous stratum, are pioneer formations that invite the colonization of vegetation on newly formed sediment banks and prepare these areas for ecological succession until a regional climax community becomes established (Veloso *et al*. 1991, IBGE 2012, Lin *et al*. 2016). The regional climax community that encompasses marshes, mangroves with herbaceous plants, and other habitats of *F. acutirostris* is the rainforest, which means that the current habitats of the species will potentially cease to be habitable. Continuous sediment accumulation and gains in altitude reduce flooding levels in estuaries, allowing plant species from more advanced stages of ecological succession to become established. Phenomena such as sediment erosion by water flow or compaction due to sediment and plant biomass weight can generate cyclical phases of regressive ecological succession involving the death of a plant community, followed by the resumption of progressive ecological succession, with the later plants adapted to new flooding conditions (M. R. Bornschein, unpubl. data).

The speed of ecological succession in the habitat of *F. acutirostris* is not well known (Reinert *et al*. 2007), but these authors reported cases of emerging herbaceous formations in two years (secondary marshes), four years (marshes between coastal dunes), and 10 years (tidal marshes); cases of emerging arboreal formations in five years (mangrove); and cases of ecological succession from marshes between coastal dunes with tree formations dominated by *Tabebuia cassinoides* and herbaceous plants within a maximum of 27 years. In this study, we report the succession from herbaceous formations to arboreal formations, with or without the presence of a lower stratum containing typical herbaceous marsh plants, within a maximum of 37–40 years, and from formations with an upper arboreal stratum and a lower stratum containing typical herbaceous marsh plants to formations without marsh species in the lower stratum within 32–40 years (Table 5). Additionally, we report cases of regressive ecological succession from tree formations without typical marsh herbaceous plants in the lower stratum (tree formation dominated by *Calophyllum brasiliense*) to the same tree formations with marsh herbaceous plants in the lower stratum (tree formation dominated by *Calophyllum brasiliense* with herbaceous plants) within a maximum of 37 years (Table 5). This may have occurred due to increased tidal height, which could have raised the height and duration of flooding and increased the salinity of the environment, killing trees and increasing light penetration into the lower stratum. This phenomenon has occurred in other parts of the world, with formations adjacent to mangroves and other estuarine formations experiencing vegetation mortality and regressive succession (Butzeck *et al*. 2016).

The conversion of tidal marshes to mangroves with herbaceous plants is not an expected path for ecological succession for either marshes or mangroves (Veloso *et al*. 1991; Reinert *et al*. 2007, IBGE 2012). Each of these environments initiates ecological succession in newly exposed sediments and succeeds to arboreal formations of trees typical of flooded areas, but at an advanced stage of succession. Therefore, it is not expected that a tidal marsh would be succeeded by arboreal formations with trees exclusive to the initial stages of ecological succession, such as mangrove trees. Mangroves are floodplain formations found in tropical regions (D’Odorico *et al*. 2012), and tidal marshes with mangrove trees are considered a particular type of marsh known as subtropical salt marsh (Bornschein *et al*. 2017). Salt marshes are floodplain formations in colder regions (D’Odorico *et al*. 2012). The replacement of a community that is characteristic of colder regions (subtropical salt marsh) by another that is characteristic of warmer regions (mangrove) can be considered a tropicalization effect (e.g., Bianchi and Morri 2003, Encarnação *et al*. 2019), possibly due to climate change.

The current data support Reinert *et al*. (2007), who stated that the greatest impact on the conservation of *F. acutirostris* is invasion by exotic grasses, which must be actively controlled and managed (Reinert *et al*. 2007, 2009, Reinert and Bornschein 2008). These grasses are widespread invasive species that have a great impact on the herbaceous environments along the southern coast of Brazil, with no barriers to their dispersion over herbaceous plants except for salinity (see above and Reinert *et al*. [2007]).

### Conservation

The assessment of the Green Status of the species demonstrated that the Conservation Legacy of conservation actions taken thus far (the eradication of exotic grasses and the establishment of conservation units) and the Conservation Dependence of the ongoing actions (the eradication of exotic grasses) are null, mainly due to the projects’ small scales. However, the Conservation Gain shows the importance of eradicating exotic grasses to maintain and increase the Green Score of the species. This action is especially effective when applied at the start of grass colonization and when the propagation rate increases (Table 6).

It is quick and relatively expensive to eradicate 1 ha of exotic grasses in estuaries where no seed bank is formed (Bornschein *et al*. 2022). Invaded tidal marshes take 10 months to recover, from the start of management to the return of native vegetation (Bornschein *et al*. 2022), although with a reduced number of species. The cost of managing 1 ha is 13,404–29,356 USD, depending mainly on the distance of the management areas from the workers’ homes (Bornschein *et al*. 2022). This means that the cost of eradicating patchy AOO of *F. acutirostris* that have been invaded and dominated by exotic grasses is 3,455,015.04–7,566,802.56 USD, without considering the extent of the plant’s spread since the assessment years (2005–2022; Table 1) and its likely spread during the years of eradication activity.

Considering the scarcity of resources for exotic plant control projects and the high costs involved, we propose the establishment of *exotic-free zones* (EFZs) as small geographic areas with a significant amount of non-invaded or little invaded environments, thus, having advantageous management costs versus benefits. This proposal is based on the premise that the protection of the species’ habitat could be maintained in the areas with the highest cost–benefit ratios, allowing the remaining areas to be considered irreversibly or functionally lost. The establishment of EFZs could be vital for territorial management, particularly within conservation units, which might include the *Roteiro Metodológico de Planejamento de Unidades de Conservação* in Brazil (Methodological Guideline for the Planning of Conservation Units; Galante *et al*. 2002).

The six proposed EFZs represent 47.4% of the area of occupation of *F. acutirostris* and 48.6% of its population (Figure 3; Table 4). We consider them strategic areas for the conservation of the species where the management of *U. arrecta* and *U. mutica* should be carried out (see details in Bornschein *et al*. [2022]). We emphasize that the available resources must be adapted to the size of the patches to be managed to ensure that the eradication of exotic species is completed without depleting resources (Moody and Mack 1988); otherwise, the plants will regrow and completely undermine the investment.

The addition of a new functional population was the main conservation action responsible for the increase in the Green Score in the Future-with-conservation scenario. Therefore, we propose the assisted colonization of the species into coastal environments within the species’ extent of occurrence to enhance conservation, such as the Acaraí river (*c.* 26°14’53”S, 48°31’59”W), located at the Acaraí State Park, in São Francisco Island, municipality of São Francisco do Sul, on northern coast of Santa Catarina, where there are 215.31 ha of similar habitats to those of the species’ occurrence with high conservation status (Figure 1). The São Francisco Island serves as an example of a location where the species possibly did not reach due to its low flight capacity, low habitat connectivity, and/or insufficient time for colonization (Reinert *et al*. 2007).

## Conclusion

*Formicivora acutirostris* is a globally threatened species due to its small population and the continuous loss of individuals caused by habitat loss due to human activities and, mainly, ecological succession. All habitats occupied by the species are pioneer formations that rapidly become established in an area, turning it into a forest no longer suitable for *F. acutirostris*. Understanding that the species’ habitat is highly dynamic and temporary is important for formulating plans for species’ conservation. Landscape transformation may limit the formation of new habitats and the ability of species to colonize them.

The invasion of exotic grasses is the most significant anthropogenic threat. It is important to continue eradicating these invasive exotic species and expanding managed areas. The costs are high, so our proposal for the establishment of EFZs indicates strategic locations where meager resources could be allocated to achieve the highest environmental returns.

The proposed assisted colonization could mitigate the loss of habitats due to ecological succession. There are suitable locations without species’ occurrence that could support introduced individuals. The largest limiting factor, as with the management of exotic species, is fundraising. A cheaper option for management action with equivalent management objectives would be the systemic management of arboreal formations by tree cutting, both of typical forest trees and mangroves, to slow and/or reverse ecological succession. However, Brazilian society and public authorities must recognize their role in preserving conservation-dependent species since, otherwise, it will be difficult to obtain authorization for such environmental management.

Possibly due to more intense and frequent flooding, regressive succession is already evident. This phenomenon, which increases the AOO of the species, may be due to climate change, and may become more frequent and widespread over time, at least along the southern coast of Brazil. However, probably it will not cause more than a minimal fraction of the loss of AOO caused by progressive ecological succession.

The advance of mangroves across tidal marshes should not be considered progressive ecological succession. This is probably another phenomenon related to climate change, known as tropicalization, which is occurring for the first time in Brazil.

Continuous monitoring of *F. acutirostris* and its habitats is strategically necessary to continually adjust management proposals and assess their effects. Gaining knowledge of this species can facilitate the implementation of conservation actions for other species facing similar challenges, both in terms of obtaining proposals and obtaining legal approval and implementation.

## Acknowledgments

Our analyses were carried out as a result of the “Olha o Clima, Litoral!” project financed by Petrobras Socioambietal. Most of the projects that yielded results for this study were developed through Mater Natura – Instituto de Estudos Ambientais, with financial management support from Helena Zarantonieli. Alexandre Bianco and João A. B. Vitto provided valuable information regarding the locations of *F. acutirostris* records in the municipality of Laguna. Claudia Golec, Ricardo Belmonte-Lopes, Felipe Shibuya, Larissa Teixeira, Maria Fernanda Ferreira Rivas, and Cecilia de Camargo Rocha assisted with many fieldwork activities. Carla S. Fontana and Márcio Repenning assisted with fieldwork in Rio Grande do Sul. Larissa Teixeira and Gabriel Marchi provided important photographs. Sérgio A. A. Morato, Mauro Pichorim, and Fernando de Camargo Passos reviewed a previous version of this work and made suggestions that improved its quality.

