## Supplementary tables S1-S5 for "Population of the Paraná Antwren *Formicivora acutirostris* reassessed as being 64% smaller: Revision of its conservation status and assessment of its Green Status, with management proposals"

Supplementary Materials

Table S1. Habitats studied along the southern coast of Brazil to estimate the density of *Formicivora acutirostris* (Dens) and evaluate its area of occupancy through analysis using aerial photographs and orbital images (Amb).

| Habitat^1^ | Classification according to Veloso *et al.* (1991) and IBGE (2012) | Study conducted | | *In loco* evaluation | Characteristics of the habitats^2^ | | |
| --- | --- | --- | --- | --- | --- | --- | --- |
|  |  | Dens | Amb |  | Characteristic herbaceous and shrubs | Arboreal stratum | Salinity |
| Tidal marsh (*brejo de maré*)^3, 4^ | Pioneer Herbaceous Formation of Fluviomarine Influence *(Formação Pioneira de Influência Fluviomarinha estágio herbáceo)* | X | X | X | *Acrostichum danaeifolium*, *Cladium mariscus*, *Crinum americanum*, *Echinodorus grandiflorus*, *Fuirena robusta*, *Schoenoplectus californicus*, *Talipariti pernambucense* (shrub), *Typha domingensis* | Absent | Present |
| Tidal marsh^3^ | Pioneer Herbaceous Formation of Fluvial Influence *(Formação Pioneira de Influência Fluvial estágio herbáceo)* |  | X | X | *Begonia fischeri*, *Commelina diffusa*, *Eleocharis interstincta*, *Eleocharis* cf. *montana*, *F. robusta*, *Hyptis* sp., *Mikania hastatocordata*, *Mikania trinervis*, *Panicum mertensii*, *Piper* sp., *Polygonum meisnerianum*, *S. californicus*, *Senna* cf. *bicapsularis* (shrub), *Stigmaphyllon ciliatum*, *Thelypteris interrupta*, *Thelypteris serrata*, *Vernonia beyrichii*, *Vigna luteola* | Absent | Absent? |
| Saw grass marsh (*brejo de capim-serra*)^3^ | Pioneer Herbaceous Formation of Fluviomarine Influence *(Formação Pioneira de Influência Fluviomarinha estágio herbáceo)* |  | X | X | *Acrostichum danaeifolium*, *C. mariscus*, *T. pernambucense* (shrub) | Absent | Present? |
| Meander marsh (*brejo de meandro*) | Pioneer Herbaceous Formation of Fluvial Influence *(Formação Pioneira de Influência Fluvial estágio herbáceo)* |  | X | X | *Eleocharis interstincta*, *Hedychium coronarium*, *T. domingensis*, *Urochloa arrecta* | Absent | Absent |
| Marsh between coastal dunes (*brejo intercordão*) | Pioneer Herbaceous Formation of Fluvial Influence *(Formação Pioneira de Influência Fluvial estágio herbáceo)* |  | X | X | *Blechnum serrulatum, C. mariscus, C. diffusa, Fuirena umbellata, Ipomoea cairica, Ipomoea purpurea, M. trinervis, Osmunda palustres, Scleria sp., Thelypteris interrupta, T. domingensis* | Absent | Absent? |
| Secondary marsh | Pioneer Herbaceous Formation of Fluvial Influence *(Formação Pioneira de Influência Fluvial estágio herbáceo)* |  | X |  | *Eleocharis interstincta*, *F. robusta*, *F. umbellata, Polygonum meisnerianum*, *Rhynchospora corymbosa*, *Scleria* sp. *T. interrupta, T. domingensis* | Absent | Absent |
| Tree formation dominated by *Calophyllum brasiliense* with herbaceous plants (*guanandizal com herbáceas*) | Pioneer Arboreal Formation of Fluviomarine Influence *(Formação Pioneira de Influência Fluviomarinha estágio arbóreo)* | X | X | X | *Acrostichum danaeifolium*, *C. mariscus*, *C. americanum*, *E. grandiflorus*, *F. robusta*, *Poa* sp., *S. californicus*, *S. ciliatum*, *T. pernambucense* (shrub) | *Annona glabra*, *Calophyllum brasiliense*, *Eugenia umbelliflora* | Present |
| Tree formation dominated by *Tabebuia cassinoides* with herbaceous plants (*caxetal com herbáceas*) | Pioneer Arboreal Formation of Fluvial Influence *(Formação Pioneira de Influência Fluvial estágio arbóreo)* | X | X | X | *Caliptrocarya longifolia*, *E. interstincta*, *Forsteronia leptocarpa*, *Oxypetalum* sp., *P. mertensii*, *P. meisnerianum*, *R. corymbosa*, *Scleria* sp. | *Annona glabra*, *Gomidesia palustres*, *Miconia jucunda*, *Myrcia multiflora*, *Rapanea* sp., *Tabebuia cassinoides* | Absent? |
| Mangrove with herbaceous plants (*manguezal com herbáceas*) | Pioneer Arboreal Formation of Fluviomarine Influence *(Formação Pioneira de Influência Fluviomarinha estágio arbóreo)* | X | X | X | *Acrostichum danaeifolium*, *C. americanum*, *S. californicus*, *T. pernambucense* (shrub) | *Laguncularia racemosa* | Present |
| Tree formation dominated by *C. brasiliense* (*guanandizal*) | Pioneer Arboreal Formation of Fluvial Influence *(Formação Pioneira de Influência Fluvial estágio arbóreo)* |  | X | X | Absent | *Annona glabra*, *C. brasiliense*, *E. umbelliflora* | Absent |
| Tree formation dominated by *T. cassinoides* (*caxetal*) | Pioneer Arboreal Formation of Fluvial Influence *(Formação Pioneira de Influência Fluvial estágio arbóreo)* |  | X | X | Absent | *Annona glabra*, *M. multiflora*, *Pouteria beaurepairei*, *Rapanea* sp., *T. cassinoides* | Absent |
| Mangrove | Pioneer Arboreal Formation of Fluviomarine Influence *(Formação Pioneira de Influência Fluviomarinha estágio arbóreo)* |  | X | X | Absent | *Avicennia schaueriana*, *L. racemosa*, *Rhizophora mangle* | Absent |
| Unidentified arboreal formation (north of 27°45’S)^2^ | Pioneer Arboreal Formation of Fluvial Influence *(Formação Pioneira de Influência Fluvial estágio arbóreo)* |  | X |  | ? | ? | Absent |
| Unidentified arboreal formation (south of 27°45’S)^2^ | Pioneer Arboreal Formation of Fluvial Influence *(Formação Pioneira de Influência Fluvial estágio arbóreo)* |  | X | X | ? | ? | Absent? |

^1^ According to Reinert *et al.* (2007), with exceptions (see below).

^2^ According to Reinert *et al.* (2007) and new data from this study.

^3^ Estuarine marshes (Doody 2001).

^4^ Subtropical salt marshes (Bornschein *et al.* 2017).

Table S2. Area of the smallest territory of a pair of *Formicivora acutirostris* for different habitats used as a reference for estimating the minimum area of habitat patches for calculating the species’ area of occupancy. A set of patches was summed to calculate the minimum area if the patches were up to 6 m apart. See the Methods section for more details.

| Habitat, according to Reinert *et al.* (2007), or a mosaic of these habitats | Minimum area estimated (ha) | Minimum area used as a reference (ha) | Source |
| --- | --- | --- | --- |
| 1- Saw grass marsh (*brejo de capim-serra*) | 3.20 |  | Reinert *et al.* (2007) |
| 2- Tidal marsh (*brejo de maré*) (with high density of *F. acutirostris*) | 0.25 |  | Reinert *et al.* (2007) |
| 3- Tidal marsh (with moderate density of *F. acutirostris*) | 0.41 |  | This study |
| 4- Tidal marsh (with reduced density of *F. acutirostris*) | 0.82 |  | This study |
| 5- Marsh between coastal dunes (*brejo intercordão*) |  | 0.41 (habitat #3) |  |
| 6- Meander marsh (*brejo de meandro*) |  | 0.41 (habitat #3) |  |
| 7- Secondary marsh |  | 0.41 (habitat #3) |  |
| 8- Tidal marsh + tree formation dominated by *Tabebuia cassinoides* with herbaceous plants (*caxetal com herbáceas*) | 0.63 |  | This study |
| 9- Tidal marsh + tree formation dominated by *Calophyllum brasiliense* with herbaceous plants (*guanandizal com herbáceas*) |  | 0.41 (habitat #3) |  |
| 10- Tidal marsh + mangrove with herbaceous plants (*manguezal com herbáceas*) |  | 0.82 (habitat #4) |  |
| 11- Tidal marsh + tree formation dominated by *C. brasiliense* with herbaceous plants + mangrove with herbaceous plants |  | 0.82 (habitat #4) |  |
| 12- Tidal marsh + tree formation dominated by *C. brasiliense* with herbaceous plants / tree formation dominated by *T. cassinoides* with herbaceous plants |  | 0.63 (habitat #8) |  |
| 13- Secondary marsh + tree formation dominated by *C. brasiliense* with herbaceous plants / tree formation dominated by *T. cassinoides* with herbaceous plants |  | 0.63 (habitat #8) |  |
| 14- Tree formation dominated by *T. cassinoides* with herbaceous plants |  | 3.20 (habitat #1) |  |
| 15- Tree formation dominated by *C. brasiliense* with herbaceous plants |  | 3.20 (habitat #1) |  |
| 16- Tree formation dominated by *C. brasiliense* with herbaceous plants (regressive ecological succession) |  | 3.20 (habitat #1) |  |
| 17- Mangrove with herbaceous plants |  | 3.20 (habitat #1) |  |
| 18- Tree formation dominated by *C. brasiliense* with herbaceous plants + mangrove with herbaceous plants |  | 3.20 (habitat #1) |  |
| 19- Tree formation dominated by *C. brasiliense* with herbaceous plants / tree formation dominated by *T. cassinoides* with herbaceous plants |  | 3.20 (habitat #1) |  |
| 20- Unidentified arboreal formation (south of 27°45’S) |  | 0.41 (habitat #3) |  |
| 21- Unidentified arboreal formation in succession of secondary marsh |  | 3.20 (habitat #1) |  |

Table S3. Variations in the density of territorial individuals of *Formicivora acutirostris* in the mosaic of tidal marsh (*brejo de maré*) with moderate density of species in a tree formation dominated by *Calophyllum brasiliense* with herbaceous plants (*guanandizal com herbáceas*) over the years at Jundiaquara Island, Guaratuba Bay, Paraná, southern Brazil.

| Year | Studied area (ha)^1^ | | Paired individuals | Density (individuals per hectare) |
| --- | --- | --- | --- | --- |
| 2006 | 11.50 | 28 | | 2.43 |
| 2007 | 11.50 | 28 | | 2.43 |
| 2008 | 11.50 | 24 | | 2.09 |
| 2009 | 11.50 | 24 | | 2.09 |
| 2010 | 11.50 | 24 | | 2.09 |
| 2011 | 11.50 | 28 | | 2.43 |
| 2012 | 11.50 | 30 | | 2.61 |
| 2013 | 11.38 | 30 | | 2.64 |
| 2014 | 11.38 | 30 | | 2.64 |
| 2015 | 11.38 | 32 | | 2.81 |
| 2016 | 11.38 | 32 | | 2.81 |
| 2017 | 11.38 | 32 | | 2.81 |
| 2018 | 11.30 | 32 | | 2.83 |
| 2019 | 11.30 | 30 | | 2.65 |
| 2020 | 11.30 | 30 | | 2.65 |
| 2021 | 11.10 | 26 | | 2.34 |
| Mean | 11.40 | 28.75 | | 2.52 |

^1^ The area varied due to erosive processes.

Table S4. Variations in the density of territorial individuals of *Formicivora acutirostris* in the mosaic of tidal marsh with moderate density of species and tree formation dominated by *C. brasiliense* with herbaceous plants over the years for Continente, Guaratuba Bay, Paraná, Southern Brazil.

| Year | Studied area (ha)^1^ | Paired individuals | Density (individuals per hectare) |
| --- | --- | --- | --- |
| 2010 | 8.26 | 20 | 2.42 |
| 2011 | 8.26 | 18 | 2.18 |
| 2012 | 8.26 | 20 | 2.42 |
| 2013 | 8.73 | 20 | 2.29 |
| 2014 | 8.73 | 20 | 2.29 |
| 2015 | 8.73 | 20 | 2.29 |
| 2016 | 8.73 | 26 | 2.98 |
| 2017 | 8.68 | 26 | 3.00 |
| 2018 | 8.68 | 24 | 2.76 |
| 2019 | 8.68 | 24 | 2.76 |
| 2020 | 8.47 | 22 | 2.60 |
| 2021 | 8.57 | 20 | 2.33 |
| Mean | 8.57 | 21.67 | 2.53 |

^1^ The area varied due to erosive processes and the spatial arrangement of certain territories. If the regularly monitored territory extended slightly beyond the study area, we expanded the study area to incorporate it. If a regularly monitored territory extended too far beyond the study area, or if a neighboring unmonitored territory extended a little further into the study area, the study area was retracted to exclude the area of these partially encompassed territories.

Table S5. Variations in the density of territorial individuals of *Formicivora acutirostris* in the mosaic of tidal marshes with reduced density of species and mangroves with herbaceous plants over the years at Folharada Island, Guaratuba Bay, Paraná, Southern Brazil.

| Year | Studied area (ha)^1^ | Paired individuals | Density (individuals per hectare) |
| --- | --- | --- | --- |
| 2009 | 16.29 | 20 | 1.23 |
| 2010 | 16.29 | 20 | 1.23 |
| 2011 | 16.29 | 26 | 1.60 |
| 2012 | 16.29 | 26 | 1.60 |
| 2013 | 16.29 | 26 | 1.60 |
| 2014 | 15.9 | 24 | 1.51 |
| 2015 | 15.9 | 24 | 1.51 |
| 2016 | 15.9 | 24 | 1.51 |
| 2017 | 15.9 | 22 | 1.38 |
| 2018 | 15.9 | 22 | 1.38 |
| 2019 | 15.8 | 20 | 1.27 |
| 2020 | 15.9 | 18 | 1.13 |
| 2021 | 16.0 | 14 | 0.88 |
| Mean | 16.05 | 22.00 | 1.37 |

^1^ The area varied due to erosive processes, changes in the extent of vegetation cover, and the spatial arrangement of certain territories. If the regularly monitored territory extended slightly beyond the study area, we expanded the study area to incorporate it.
